# Spatially unitized associations occupy an intermediate position along the item-to-association continuum in older adults

**DOI:** 10.64898/2026.07.17.739184

**Authors:** Min Sung Seo, Alexa Becker, Lin-Han Hannah Huang, Amy A. Overman, Nancy A. Dennis

## Abstract

Older adults show disproportionate impairments in associative memory relative to memory for individual items. Unitization, the process by which discrete elements are encoded as a single integrated representation, has been proposed to mitigate this deficit by reducing reliance on hippocampal binding and shifting processing toward perirhinal cortex-mediated item-based encoding. However, most prior work has not included a single item baseline, leaving unclear whether unitized associations are truly processed like items. Here, we investigated whether spatial proximity supports associative memory in older adults and how this benefit is reflected in medial temporal lobe (MTL) encoding activity. Participants encoded unrelated object pairs presented either proximally (spatially integrated) or distally (spatially separated), alongside a single item condition. Behaviorally, older adults showed better memory for proximal than distal pairs, though performance remained below that of single items. Neurally, MTL activity scaled with associative demand, with hippocampal activity increasing and perirhinal activity decreasing as demand increased. Results further revealed an anterior-posterior MTL dissociation, with proximal pairs sharing neural properties with both single items and distal pairs without uniquely recruiting either. Furthermore, hippocampal patterns reliably distinguished item from associative encoding, and the parahippocampal cortex discriminated all three conditions, indicating that the item-associative structure of the MTL was preserved in aging. Together, these findings suggest that proximal pairs occupy an intermediate position between item and associative representations within an intact MTL architecture, and that spatial proximity benefits associative memory in older adults by partially, but not fully, shifting associative demands toward item-based processing.

**Highlights:**

- Spatial proximity improved associative memory in older adults.
- Hippocampal and PRC activity scaled with associative binding demands.
- Spatially unitized associations fell between single items and distal associations in MTL encoding activity.
- MTL item-association functional structure was preserved in older adults.

## 1 Introduction

Associative memory refers to the ability to form and retrieve links between discrete elements of an event, such as those among items or between items and their contextual features (Campbell et al., 2010; Chalfonte & Johnson, 1996; Naveh-Benjamin, 2000; Spencer & Raz, 1995). A well-established finding in the cognitive aging literature is that older adults are disproportionately impaired in associative memory relative to memory for individual items (Naveh-Benjamin & Mayr, 2018; Old & Naveh-Benjamin, 2008a). This deficit has been attributed to age-related declines in the ability to bind item-item and item-context relationships during encoding and the subsequent retrieval of these bindings at test (Naveh-Benjamin, 2000; Naveh-Benjamin et al., 2004; Old & Naveh-Benjamin, 2008b). Older adults have been shown to exhibit poorer associative memory performance compared to younger adults across a wide range of stimuli, including word pairs (De Brigard et al., 2020; Kim & Giovanello, 2011), image pairs (Guez & Lev, 2016), face-scene (Dennis et al., 2008), and face-name associations (Naveh-Benjamin et al., 2004, 2009).

Given these findings, researchers have sought to identify mechanisms that may mitigate associative memory deficits in aging. One such mechanism is unitization, defined as the process by which two or more distinct elements are encoded as a single unified representation (Bastin et al., 2013; D’Angelo et al., 2017; Graf & Schacter, 1989). By transforming multiple elements into a unified representation, unitization is thought to reduce the need for effortful associative binding and instead shift memory processing toward item-based mechanisms, which are relatively preserved with age (Dennis et al., 2024; Diana et al., 2008; Parks & Yonelinas, 2015). Consistent with this account, unitization has been shown to enhance associative memory across a range of paradigms and stimulus types, including word pairs (Haskins et al., 2008; Quamme et al., 2007; Zheng et al., 2015), image pairs (Bridger et al., 2017; Dennis et al., 2025; Huffer et al., 2022), word-color (Bastin et al., 2013; Zheng et al., 2016), and face-occupation associations (Overman & Stephens, 2013; Ricupero et al., 2023). For example, perceptual or visual integration has been used to induce unitization by presenting unrelated objects in a meaningful or spatially coherent configuration (e.g., an airplane positioned above a telescope; an otter holding a cleaver), encouraging participants to encode the pair as a single, integrated image (Bridger et al., 2017; Dennis et al., 2025; Huffer et al., 2022). Similarly, semantic integration has been achieved by combining unrelated words into novel compound concepts (e.g., “slope-bread”) through a unifying definition (“A pastry eaten by mountain-climbers”), promoting the encoding of the pair as single conceptual unit rather than two independent words (Bader et al., 2010; Haskins et al., 2008; Quamme et al., 2007). In both cases, experimental manipulations are intended to facilitate the formation of integrated representations that reduce reliance on effortful associative binding, with converging findings showing enhanced memory for unitized stimuli, relative to non-unitized stimuli.

Neuroimaging studies using functional magnetic resonance imaging (fMRI) have sought to characterize the neural mechanisms underlying unitization, particularly within medial temporal lobe (MTL) regions. Much of this work has been interpreted within the Binding of Items and Context (BIC) model, which posits that the MTL distinguishes between item-based and associative memory processes, with the perirhinal cortex (PRC) supporting item-level and familiarity-based representations, the parahippocampal cortex (PHC) supporting contextual and spatial representations, and the hippocampus supporting the binding of multiple elements into relational memory representations (Diana et al., 2007; Ranganath, 2010). Accordingly, if two items are successfully unitized such that they are encoded as a single integrated representation, the BIC model predicts a shift away from hippocampal-dependent associative binding and toward PRC-mediated item-based processing.

However, neuroimaging evidence for this predicted shift is mixed. On one hand, patients with hippocampal damage, who are impaired in recollection but relatively spared in familiarity, show preserved or even enhanced associative recognition for unitized word pairs relative to non-unitized pairs, suggesting that unitization does reduce reliance on hippocampal binding processes (Giovanello et al., 2006; Quamme et al., 2007). Similarly, fMRI studies using preexisting or strongly integrated associations (e.g., compound words or logically integrated objects) have reported greater PRC involvement during encoding of unitized stimuli, alongside increased hippocampal engagement for non-unitized associations (Ford et al., 2010; Memel & Ryan, 2017). One study found increased PRC activation during encoding and retrieval of unrelated word pairs that were integrated into novel compound concepts via definitions (“slope-bread” being “A pastry eaten by mountain-climbers”), with PRC activity predicting subsequent familiarity-based recognition.

On the other hand, other studies have reported mixed patterns of activation across MTL regions, with findings showing continued involvement of the hippocampus and PHC even under conditions intended to promote unitization (Bader et al., 2014; Dennis et al., 2019, 2025; Memel & Ryan, 2017; Ricupero et al., 2023). For example, using a similar definition-based paradigm to induce unitization, Bader et al. (2014) did not observe increased PRC involvement for unitized associations. Instead, unitization was associated with reduced engagement of the recollection network and greater recruitment of regions including the PHC and medial prefrontal cortex during retrieval. Taken together, these findings indicate that unitization is not a uniform process, but rather varies to the extent to which associations are successfully transformed into item-like representations, with corresponding variability in the balance of PRC, PHC, and hippocampus engagement (Parks & Yonelinas, 2015).

Critically, despite the central claim that unitization promotes more item-like processing of associative information, most prior work has evaluated this hypothesis indirectly by comparing unitized and non-unitized associations without including a single item baseline. As a result, it remains unclear whether unitized representations are processed similarly to individual items, or instead reflect an intermediate representation that continues to rely, at least in part, on associative binding mechanisms. This limitation is particularly important in the context of the BIC model, which links item-level processing to PRC function and associative binding to hippocampal engagement (Diana et al., 2007; Ranganath, 2010). Including a single item condition therefore provides a more direct test of the extent to which unitization shifts processing toward item-based representations. If unitized associations more closely resemble item representations, they should elicit neural responses more similar to single items, particularly within PRC. Alternatively, if unitized associations continue to rely on associative binding mechanisms, their neural activity should more closely resemble that of non-unitized associations, including greater hippocampal involvement.

Building on this question, prior behavioral work from our lab has directly examined whether unitized associations function similarly to single items by including an explicit single item baseline during memory retrieval. Using a recognition memory paradigm, Carpenter & Dennis (2023) demonstrated that although unitized pairs showed improved recognition relative to associative pairs across both younger and older adults, their performance did not reach the level of single item memory. Instead, memory for unitized word pairs appeared to fall between that of item and non-unitized associative memory, consistent with the idea that unitization may enhance associative binding without fully transforming representations into true item-like forms. Extending this behavioral work to neuroimaging, recent fMRI work from our lab examined these questions in younger adults using the same paradigm employed in the current study (Dennis et al., 2025). Specifically, two associative memory conditions were investigated alongside a single object condition. In the proximal associative condition, unrelated object pairs were presented in an integrative and logical orientation, in order to encourage integrated or unitized processing, while in the distal associative condition, unrelated object pairs were presented at a set distance in order to promote more typical associative binding between distinct items. Behaviorally, proximal pairs produced better recognition memory than distal pairs, indicating that proximal placement reduced the demands of associative memory, although performance remained below that of single items. Neurally, results revealed that, despite their behavioral benefit, within subregions of the MTL, proximal pairs exhibited neural responses more similar in magnitude and location to that of distal pairs than single items. Specifically, compared to single items, both distal and proximal pairs showed greater activation in the bilateral PHC at both encoding and retrieval. However, distal pairs exhibited greater PHC recruitment than both proximal pairs and single items. This suggests that the proximal image pairs may have still required item-item binding resources for successful memory, albeit less so than that required by distal pairs. Additionally, multivariate pattern similarity analysis (PSA) found that voxel-level neural representations of proximal pairs were more similar to that of distal pairs than single items within both the MTL and cortical associative memory network regions including the prefrontal cortex, superior parietal lobule and precuneus. Together, these findings suggest that proximal pairs appeared to occupy an intermediate position between single item and associative representations, retaining characteristics of associative processing while requiring fewer binding-related resources than distal pairs.

The current study extended this work to older adults and tested whether proximal placement similarly benefits associative memory in aging, and where these representations fall along the continuum between item-based and associative processing with respect to MTL activation. We focused on older adults specifically, because associative memory is disproportionately impaired in aging (Naveh-Benjamin, 2000; Old & Naveh-Benjamin, 2008a), making them the population for whom unitization-based support is most consequential, and because the MTL processing of item and associative memory has been relatively less investigated in older adults (Dennis et al., 2024). Whether this functional organization is maintained in older adults, and how unitized associations are situated within it, remains an open question. Rather than testing age differences directly, which our companion study in younger adults addresses (Dennis et al., 2025), the present study examined how the aging MTL supports proximity-based associative memory, and where proximal representations fall relative to item and associative processing within the older adult MTL.

Using the same recognition paradigm from Dennis et al. (2025), unrelated object pairs were presented either proximally, or distally, alongside a single item condition. Critically, the inclusion of single, proximal, distal conditions allowed us to directly characterize the extent to which unitized (“proximal”) pairs process in an item-based or associative manner within the MTL. Specifically, using both univariate and multivariate fMRI approaches, we examined whether neural activity associated with proximal object pairs more closely resembled item-based or associative representations within MTL regions. Based on prior work and the BIC framework, we expected MTL activity to vary as a function of associative binding demand, with a) greater hippocampal involvement associated with conditions requiring greater binding and b) greater PRC involvement associated with more item-based processing.

## 2 Methods

### 2.1 Transparency and openness

Preregistration of the hypotheses, methods, analyses, stimulus materials, de-identified data, and analyses scripts are publicly available on the Open Science Framework (OSF: 10.17605/OSF.IO/3D75J).

### 2.2 Participants

Participants were recruited from greater Centre County, PA. All participants were screened for psychiatric and neurological disorders, head injury, stroke, learning disability, medication that affects cognitive and neural function, and substance abuse. Furthermore, participants were right-handed, had normal (or corrected-to normal) vision, and were either native English speakers or had first learned English before the age of six. Power analysis conducted through the base *power.t.test* function in R revealed that 30 participants would be required to achieve 75% power at a two-sided 5% alpha level to identify a medium within-person effect (*d_z_* = 0.5). Written informed consents approved by The Pennsylvania State University institutional review board were provided to the participants before the testing.

Thirty-seven participants were tested, with 2 removed from analyses due to discomfort in the MRI scanner (*n* = 1) and response rates below 90% (*n* = 1), resulting in a final sample of 35 participants (ages 60–85 years; *M* = 70.9, *SD* = 5.25; 26 females, 9 males). This sample provided 81.95% power to detect a within-subject effect of *d_z_* = 0.5. Participants identified as White (*n* = 32), Asian (*n* = 1), Black or African American (*n* = 1), or more than one race (n = 1).

### 2.3 Materials and procedure

Materials and procedure were identical to those reported previously in Dennis et al. (2025). For completeness and ease of reference, the details are reproduced below.

The experiment consisted of three conditions: single item memory and two associative memory conditions, proximal and distal (described below). Stimuli were images taken from The Bank of Standardized Stimuli (BOSS) database (Brodeur et al., 2010). Images were pseudo-randomly selected from the database and associative pairs were pseudo-randomly assigned, such that we ensured equitable numbers of animate to inanimate objects across our three conditions. Animate objects were also never paired with one another. Associative pairs were then normed (across 5 younger adults) to eliminate any pre-experimental semantic associations between paired images. For each pairing, participants rated on a scale of 1-5 how semantically related the two images were (1 = extremely unrelated; 5 = extremely related). To ensure they understood the rating scale, participants were first given examples of pairings that have a rating of 1 (a bird and a clipboard, since they aren’t usually related in any way), 3 (a bird and a butterfly, since they are both animals), and 5 (a bird and a nest, since a bird can live in a nest). Only image pairings that had an average rating lower than 3 were used in the study. The logicality of image orientation for the proximal pairings was then normed by second set of individuals (N=23). First, the image pairings that were determined to be semantically unrelated by the earlier semantic norming process were oriented in a way deemed illogical by the researchers. The logicality norming participants then rated on a scale of 1-5 how logical the orientation of each pairing was (1 = not logical; 5 = logical). Though all image pairings consisted of unrelated items, and thus their simple cooccurrence in the real world would be illogical, participants were instructed to put this fact aside and indicate what the best orientation would be if the two items did occur together. To ensure they understood the rating scale, participants were first given examples of image orientations that would have a rating of 1 (a roll of aluminum foil placed on top of a bird’s head) and 5 (a bird placed atop a roll of aluminum foil, as if it were perched on it). If a participant scored any pairing below a 3, they were asked to explain what they felt would be a more logical orientation of the images. Any image pairs rated a 3 or lower were oriented in a different manner by the researchers, based on participant feedback, and then sent through another round of rating with different raters. This entailed six rounds of logicality norming (round 1 n = 5; round 2 n = 3; round 3 n = 3; round 4 n = 4; round 5 n = 4; round 6 n = 4) until all image pairings had an average rating greater than 3. These pairings were used in the present study. To control for any differences between image pairs in each condition, image pairs were counterbalanced across two tasks, in which the pairings that were assigned to the proximal condition in version A were assigned to the distal condition in version B, and vice versa. There were no significant differences between counterbalancing versions, and as such all results are collapsed across versions.

The encoding phase occurred in the scanner and consisted of 144 trials equally split across 4 runs, including 48 distal trials, 48 proximal trials, 28 single trials, and 20 randomly distributed attention check trials. We included a greater number of distal and proximal trials to account for the recombination of pairs at retrieval as lures, and also in order to have the same number of single, distal and proximal lures at retrieval. All images were presented at a distance of approximately 133.5 cm. In the distal condition, the two unrelated images were presented at a distance of six inches apart (6° 32’ 0.02’) apart from one another (9° 47’ 0.23’ total width). In the proximal condition, the two unrelated images were presented in the center of the screen (6° 32’ 0.02’ width), slightly overlapping and oriented in a logical manner (see Figure 1). In the single condition, one image was presented in the center of the screen (3° 16’ 0.17’ width). The experiment was presented using E-Prime (Psychology Software Tools, Inc., 2016).

**Figure 1.**
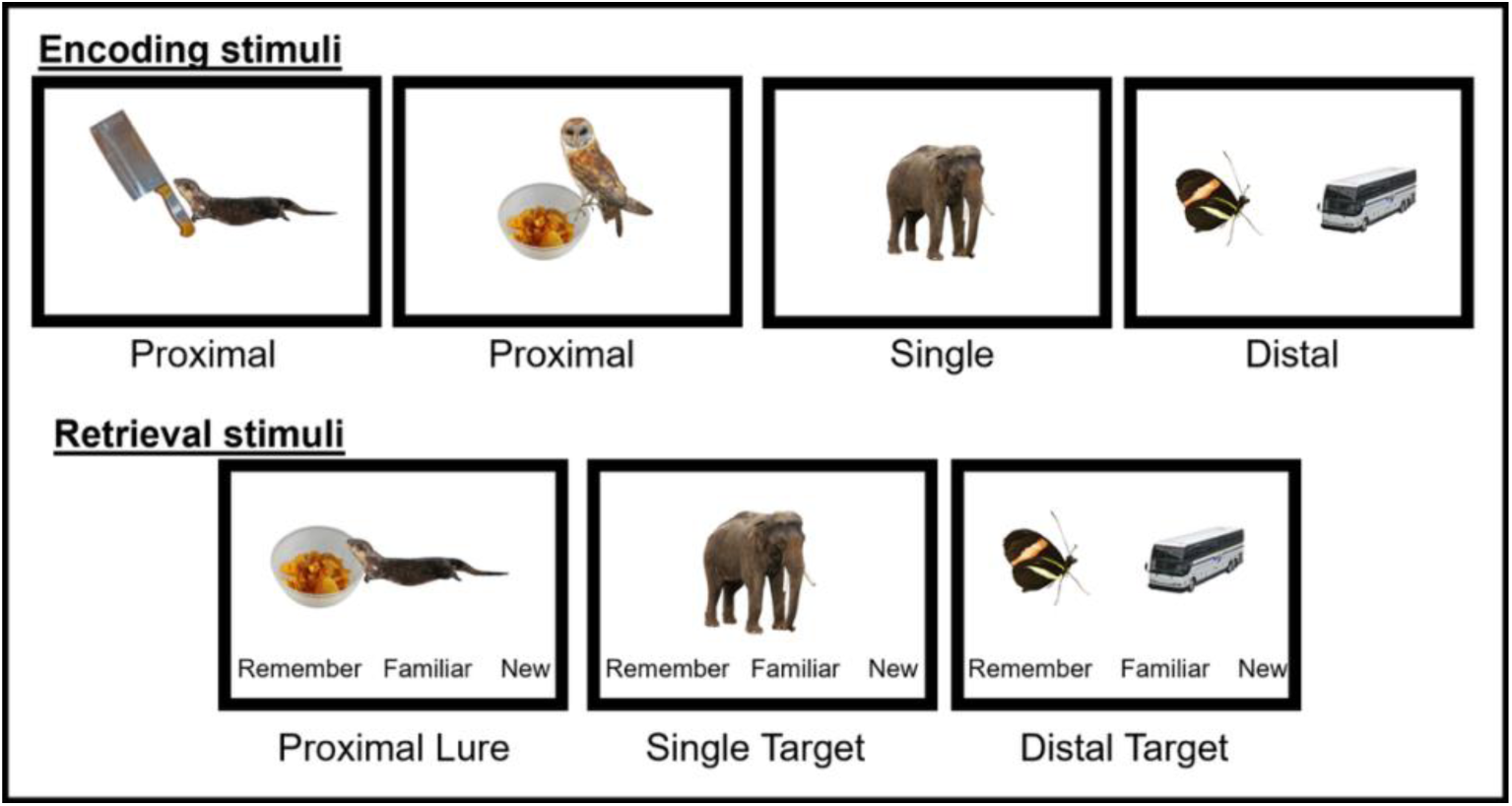
Stimuli. Examples of proximal, single, and distal encoding trials, as well as target and recombined lure trials. *Adapted from Dennis et al. (2025)*.

Encoding trials were presented for 3000 ms each and separated by a jittered interstimulus interval lasting 1500, 3000, or 5000 ms at random. Participants were asked to study all of the images and try to remember them for a later memory test. As an attention check, participants were instructed to respond with their index finger every time they saw a watermelon, which was presented both individually and in pairs. After the encoding phase, participants were given a break in the scanner during which they viewed a silent video of fireworks while a structural scan was taken. The retrieval phase consisted of 144 trials equally split across 4 runs, including 48 distal trials, 48 proximal trials, and 48 single trials. Retrieval trials were presented for 4000 ms each and separated by a jittered interstimulus interval lasting 1500, 3000, or 5000 ms at random. Across all 4 runs of encoding, within each condition, 28 trials were subsequent targets and 20 were subsequent lures. As the current project was primarily interested in true memory, we included more target trials than lures to allow for more power in our hypotheses of interest. Target images and image pairs were presented exactly as they were during the encoding phase. In the distal and proximal conditions, lures were created by recombining images from a subset of image pairs presented during encoding to create novel pairings (see Figure 1). Recombined pairs from the associative conditions were only recombined within condition (i.e., an image that was part of a proximal pairing during encoding was never used as part of a distal lure at retrieval). In the single condition, lures were novel single images.

During retrieval, which also took place in the scanner, participants completed a modified version of the “Remember-Know-New” response paradigm, with instructions focused on confidence. In line with past research aligning recollection and familiarity with levels of confidence in memory responses (e.g., Otten, 2007; Wais et al., 2008; Wixted & Mickes, 2010; Wixted & Stretch, 2004; Yonelinas, 2002) participants were instructed to respond “Remember” with their index finger if they remembered the image or image pair to be exactly the same as what they saw in the encoding phase. They were instructed to respond “Familiar” with their middle finger if the image or image pairing was familiar to them such that they believed it was presented previously but were not completely sure. They were instructed to respond “New” if they did not remember the image or image pair from the encoding phase and believed that it was a new image or image pair. Additionally, the following text was provided on the computer screen preceding each retrieval run: “In this task, you will again see images. Some of the images and image pairs are exactly the same as before, meaning you have seen them during the study task. However, some images are new and also, some of the images that originally appeared in a pair have now been rearranged into new pairings. For image pairs please only respond that a pair is old if you saw the pair of images presented together in the study phase. Again, you are asked to respond to what images and image pairs are THE SAME as what you saw previously. Please respond with the following choices. Respond “Remember” if you remember the image or image pair to be exactly the same as you saw in the first part of the study. Respond “Familiar” if the image or image pairing is familiar to you such that you believe it was presented previously, but you are not completely sure. Respond “New” if you do not remember the image or image pair from the first part of the study and believe that it is a new image or image pairs.”

Prior to entering the scanner, participants first gave informed consent and filled out a demographic survey. The experimenter then explained the instructions and response options to them, and they completed 9 encoding and 7 retrieval practice trials to ensure they understood the nature of the task. Participants then went into the scanner to complete the task. The same instructions presented during the encoding practice were re-presented to participants at the start of encoding and in between each run of encoding. The entire encoding phase lasted approximately 20 minutes. After the encoding phase, participants were given a break in the scanner during which they viewed a silent video of fireworks while a structural scan was taken. Participants then began the retrieval phase. The same instructions presented during the retrieval practice were re-presented to participants at the start of retrieval and in between each run of retrieval. The entire retrieval phase lasted approximately 17 minutes. Responses were collected using a 4-button MR-compatible button box. At the end of retrieval, participants remained in the scanner for 1 more minute while field mapping was run. Participants then exited the scanner and were debriefed on the goal of the study and paid for their time.

### 2.4 Image acquisition

Structural and functional images were acquired using a Siemens 3-T scanner equipped with a 20-channel head coil, parallel to the AC-PC plane. Structural images were acquired with a 1700-msec repetition time, a 2.280-msec echo time, a 256-mm field of view, 192 axial slices, and a 1.0-mm slice thickness for each participant. Echoplanar functional images were acquired using a descending acquisition, a 2500-msec repetition time, a 25-msec echo time, a 240-mm field of view, a 90° flip angle, and 42 axial slices with a 3.0-mm slice thickness resulting in 3.0-mm isotropic voxels.

#### 2.4.1 fMRIPrep

The following text is boilerplate text resulting from preprocessing in fMRIPrep 20.1.1 (Esteban et al., 2019; RRID:SCR_016216), which is based on Nipype 1.5.0 (Esteban et al., 2022; Gorgolewski et al., 2011; RRID:SCR_002502).

#### 2.4.2 Anatomical Data Preprocessing

The T1-weighted (T1w) image was corrected for intensity non-uniformity with *N4BiasFieldCorrection* (Tustison et al., 2010), distributed with ANTs 2.2.0 (Avants, Epstein, Grossman, & Gee, 2008, RRID:SCR_004757), and used as T1w-reference throughout the workflow. The T1w-reference was then skull-stripped with a Nipype implementation of the *antsBrainExtraction.sh* workflow (from ANTs), using OASIS30ANTs as target template. Brain tissue segmentation of cerebrospinal fluid, white matter, and gray matter was performed on the brain-extracted T1w using FAST (FMRIB Software Library 5.0.9, RRID:SCR_002823; Zhang, Brady, & Smith, 2001). Brain surfaces were reconstructed using *recon-all* (FreeSurfer 6.0.1, RRID:SCR_001847; Dale, Fischl, & Sereno, 1999), and the brain mask estimated previously was refined with a custom variation of the method to reconcile ANTs-derived and FreeSurfer-derived segmentations of the cortical gray matter of Mindboggle (RRID:SCR_002438; Klein et al., 2017). Volume-based spatial normalization to one standard space (MNI152NLin2009cAsym) was performed through nonlinear registration with antsRegistration (ANTs 2.2.0), using brain-extracted versions of both T1w reference and the T1w template. The following template was selected for spatial normalization: ICBM 152 Nonlinear Asymmetrical template version 2009c (Fonov, Evans, McKinstry, Almli, & Collins, 2009; RRID:SCR_008796; TemplateFlow ID: MNI152NLin2009cAsym).

#### 2.4.3 Functional Data Preprocessing

For each of the eight BOLD runs found per subject (across all tasks and sessions), the following preprocessing was performed. First, a reference volume and its skull-stripped version were generated using a custom methodology of fMRIPrep. Head-motion parameters with respect to the BOLD reference (transformation matrices, and six corresponding rotation and translation parameters) are estimated before any spatiotemporal filtering using MCFLIRT (FMRIB Software Library 5.0.9; Jenkinson, Bannister, Brady, & Smith, 2002). Susceptibility distortion correction was omitted. The BOLD reference was then coregistered to the T1w reference using bbregister (FreeSurfer), which implements boundary-based registration (Greve & Fischl, 2009). Coregistration was configured with six degrees of freedom. The BOLD time-series (including slice-timing correction when applied) were resampled onto their original, native space by applying the transforms to correct for head motion. These resampled BOLD time-series will be referred to as *preprocessed BOLD in original space*, or just *preprocessed BOLD*. The BOLD time-series were resampled into standard space, generating a *preprocessed BOLD* run in *MNI152NLin2009cAsym space*. First, a reference volume and its skull-stripped version were generated using a custom methodology of fMRIPrep. Several confounding time-series were calculated based on the *preprocessed BOLD*: framewise displacement (FD), derivative of root mean squared variance over voxels (DVARS), and three region-wise global signals. FD was computed using two formulations following Power (absolute sum of relative motions, Power et al., 2014; and Jenkinson [relative root-mean-square displacement between affines], Jenkinson et al., 2002). FD and DVARS are calculated for each functional run, both using their implementations in Nipype (following the definitions by Power et al., 2014). In addition, a set of physiological regressors were extracted to allow for component-based noise correction (CompCor; Behzadi, Restom, Liau, & Liu, 2007). The head-motion estimates calculated in the correction step were also placed within the corresponding confounds file. Frames that exceeded a threshold of 0.5-mm FD or 1.5 standardized DVARS were annotated as motion outliers. All resamplings can be performed with a single interpolation step by composing all the pertinent transformations (i.e., head-motion transform matrices, susceptibility distortion correction when available, and coregistrations to anatomical and output spaces). Gridded (volumetric) resamplings were performed using *antsApplyTransforms* (ANTs), configured with Lanczos interpolation to minimize the smoothing effects of other kernels (Lanczos, 1964). Nongridded (surface) resamplings were performed using *mri_vol2surf* (FreeSurfer).

Many internal operations of fMRIPrep use Nilearn 0.6.2 (Abraham et al., 2014, RRID:SCR_001362), mostly within the functional processing workflow. For more details of the pipeline, see the section corresponding to workflows in fMRIPrep’s documentation.

#### 2.4.4 Copyright Waiver

The above boilerplate text was automatically generated by fMRIPrep with the express intention that users should copy and paste this text into their article unchanged. It is released under the CC0 license.

### 2.5 Behavioral data analysis

Recollection-based discriminability (recollected d’) was analyzed using a linear mixed-effects model with condition (single, proximal, distal) as a fixed effect and subject as a random intercept. Statistical significance was assessed using Type II Wald chi-square tests. Pairwise comparisons between conditions were conducted using estimated marginal means with Benjamini-Hochberg correction for multiple comparisons (Benjamini & Hochberg, 1995).

### 2.6 fMRI analyses

#### 2.6.1 Anatomical ROIs

All analyses were conducted within anatomical MTL ROIs including the hippocampus, PRC, and PHC. ROIs were derived from a combined MTL mask, with the PRC taken from Holdstock et al. (2009), and the hippocampus and PHC taken from the Wake Forest University’s AAL PickAtlas (Maldjian et al., 2003). This same set of anatomical ROIs was used for both univariate and multivariate analyses. For univariate analyses, statistical maps were thresholded at an uncorrected threshold of *p* < .005 and cluster-level corrected to *p* < .05, using 3dClustSim in AFNI Version 21.3.04 (Cox, 1996; Cox & Hyde, 1997).

#### 2.6.2 fMRI univariate: parametric modulations

To test whether MTL activity varied as a function of associative binding demands, we conducted three separate parametric modulation analyses, modeling encoding conditions along a continuum from item-based to associative processing within the general linear model (GLM) framework in SPM12 (Ashburner et al., 2014). At the first level, trial onsets were modeled as stick functions convolved with the canonical hemodynamic response function. A single regressor capturing all recollected hit trials, (i.e., targets receiving a “Remember” response) was included and modulated by condition-specific weights reflecting associative demand. Three alternative coding schemes were specified to test competing theoretical assumptions about the position of proximal pairs along the item-to-association continuum: (1) a linear-spacing model (1:3:5), in which proximal pairs (parametric weighting of a 3) were treated as falling equidistant between single item and distal conditions (parametric weighting of 1 and 5 respectively); (2) a single biased model (1:2:5), in which proximal pairs were modeled as more similar to single items (proximal parametric weighting of a 2), consistent with strong unitization; and (3) an association-biased model (1:4:5), in which proximal pairs were modeled as more similar to distal pairs (proximal parametric weighting of a 4), consistent with heavier reliance on associative binding. Parametric modulators were mean-centered and entered into the GLM with first-order polynomial expansion. Parameter estimates from first-level models were entered into second-level random-effects analyses, and one-sample *t*-tests were conducted to identify regions showing significant positive or negative scaling with increasing associative demand. Additional regressors included trials receiving “Know” or “New” responses, all lure trials, and six head motion parameters, all of which were treated as regressors of no interest.

#### 2.6.3 fMRI univariate: between-condition contrasts

To complement the parametric modulation analyses examining scaling demands related to encoding of proximally configured associative pairs, we conducted a separate GLM using direct between-condition contrasts to further characterize differences in MTL activity across encoding conditions. At the first level, separate regressors were specified for recollected hit trials (i.e., targets receiving a “Remember” response) in the single, proximal, and distal conditions. Additional regressors included trials receiving “Know” and “New” responses, all lure trials, and six head-motion parameters, all of which were treated as regressors of no interest. First level contrast images were generated for pairwise comparisons between all three encoding conditions. Contrast images were then entered into second level random-effects analyses using one-sample t-tests to examine differences in MTL recruitment across encoding conditions. Analyses again focused on recollected target trials to allow direct comparison of neural activity associated with item-based (single), unitized (proximal), and associative (distal) encoding conditions. Based on the unitization account and the BIC framework, we expected greater hippocampal involvement for distal relative to single conditions, reflecting increased associative binding demands, and greater PRC involvement for single relative to distal conditions, reflecting greater item-based processing. Proximal pairs were expected to exhibit activity intermediate between single and distal conditions.

#### 2.6.4 fMRI multivariate: MVPA classifier

Finally, we conducted multivoxel pattern analysis (MVPA) to assess the discriminability of encoding conditions within MTL regions. Specifically, we tested whether neural patterns associated with proximal targets could be reliably distinguished from single and distal targets, providing additional tests of whether unitized representations were distinct from item and associative representations. Separate GLMs were fit in SPM using normalized unsmoothed data, with one regressor defined for each trial at encoding (Op de Beeck, 2010; Zeithamova et al., 2017). Six head motion parameters were included in each run as nuisance regressors. This approach yielded whole-brain beta parameter maps for each trial and participant. Within each parameter map, voxel values represented trial-specific beta estimates derived from a multiple regression model that included regressors for all other trials in the run as well as motion parameters.

Trial-wise beta maps were concatenated across encoding runs and submitted to classification analyses using the CoSMoMVPA toolbox (Oosterhof et al., 2016). A linear support vector machine (SVM) classifier was applied within each ROI using all voxels. Rather than a single three-class classifier, pairwise binary classifications were conducted between conditions (single vs. distal, single vs. proximal, and proximal vs. distal targets). This approach was chosen deliberately to assess whether neural representations of proximal pairs more closely resembled single or distal conditions, a question that a three-class classifier would not directly address, as it only indicates whether the three conditions are discriminable overall. A leave-one-run-out cross-validation procedure was used, in which the classifier was trained on three runs and tested on the held-out run, with classification accuracy averaged across folds to obtain subject-level estimates. Medial temporal lobe ROIs included the hippocampus, PRC and PHC.

To determine whether classification accuracy exceeded chance (50%), one-tailed one-sample *t*-tests were conducted within each ROI, with false discovery rate (FDR) correction applied across ROIs using the Benjamini-Hochberg procedure (Benjamini & Hochberg, 1995). To further evaluate classifier performance, permutation testing (10,000 iterations) was conducted by randomly shuffling trial labels within runs while preserving run structure. Across participants and ROIs, null distributions were centered near chance (mean ≈ 0.50), supporting the interpretation that observed classification accuracies reflected condition-related neural patterns rather than methodological artifacts.

## 3 Results

### 3.1 Behavioral results

Recollected d’ showed a significant main effect of condition, χ²(2) = 333.25, *p* < .001 (see Figure 2). Pairwise comparisons revealed that recollected d’ was significantly higher for single relative to both proximal (β = 1.60, 95% CI [1.36, 1.84], *t*(68) = 13.22, *p* < .001) and distal conditions (β = 2.12, CI [1.88, 2.37], *t*(68) = 17.51, *pv* < .001). Critically, proximal recollected d’ was also significantly higher than distal (β = 0.52, CI [0.28, 0.76], *t*(68) = 4.30, *p* < .001), indicating that older adults showed a memory advantage for proximal relative to distal pairs. Of note, analyses using overall d’ (using targets receiving “Remember” or “Know” responses) yielded the same pattern of results, showing significantly greater memory performance for single, followed by proximal, and then distal conditions (see Supplementary Materials).

**Figure 2.**
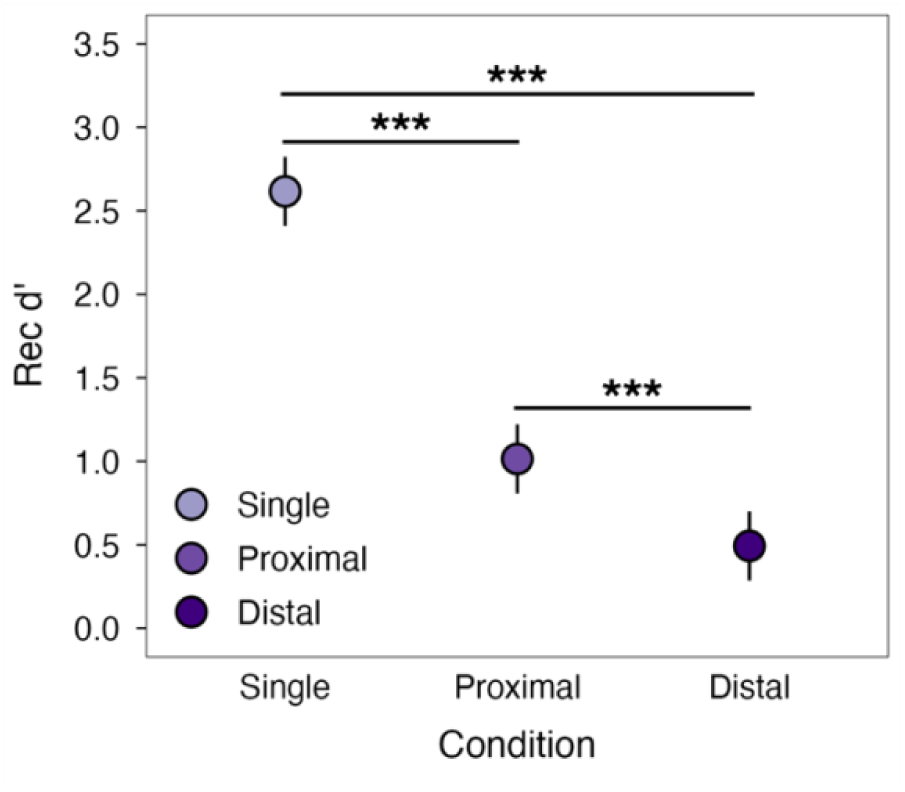
Recollected d’ across encoding conditions. Memory performance was highest for single, followed by proximal, and distal conditions. Points represent model estimated marginal means; error bars represent 95% confidence intervals; asterisks indicate significant pairwise comparisons (***p < .001).

### 3.2 fMRI results

#### 3.2.1 Parametric modulations

Parametric modulation analyses examined whether activity during encoding scaled with increasing associative demands under three alternative coding schemes (1:3:5, 1:2:5, and 1:4:5) where 1 and 5 always represented single and distal conditions, respectively, and the proximal condition varied accordingly (see Table 1 and Figure 3). Under the linear-spacing model (1:3:5), increasing associative demand was associated with greater activity in the right posterior hippocampus, alongside decreases in bilateral anterior MTL regions including the PRC. A similar pattern was observed under the single-biased model (1:2:5), with stronger increases in the right posterior hippocampus extending into PHC, and decreases in bilateral anterior MTL regions including the PRC, as well as a smaller cluster extending into anterior temporal cortex. In contrast, the association-biased model (1:4:5) did not yield any regions showing increased activity with associative demand. Instead, this model was characterized by widespread decreases in anterior MTL regions, including bilateral PRC, extending into the right anterior hippocampus.

**Figure 3.**
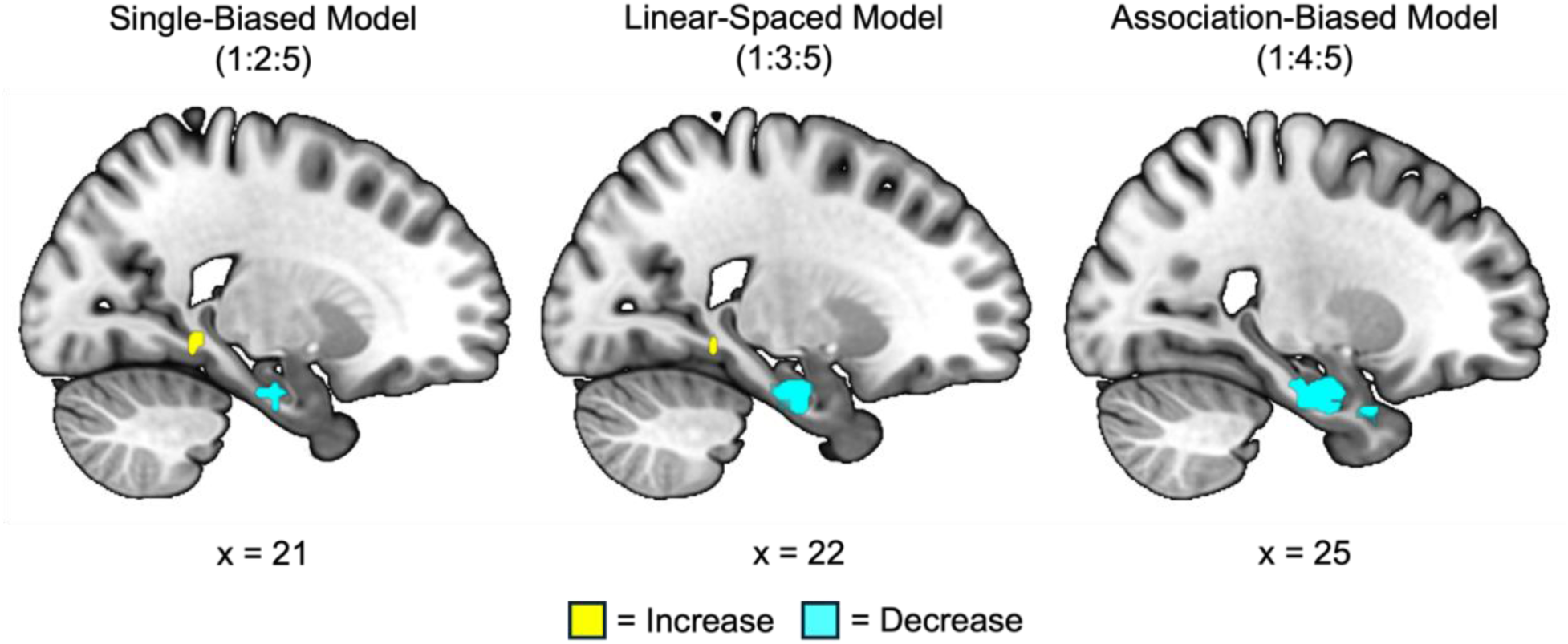
Parametric modulation of MTL activity under three coding schemes. Sagittal slices show clusters where activity scaled positively (colored yellow) or negatively (colored blue) with increasing associative demand, where single item and distal pairs were weighted 1 and 5, respectively, and the proximal weighting varied across models.

**Table 1.** Significant clusters from parametric modulation analyses of recollected hits across single, proximal, distal trials, within the MTL.

| Coding<br>(single:proximal:distal) | Regions | H | *k* | *t* | MNI Coordinates |  |  |
| --- | --- | --- | --- | --- | --- | --- | --- |
|  |  |  |  |  | *x* | *y* | *z* |
| 1:3:5 |  |  |  |  |  |  |  |
| Increase | Posterior HC | R | 11 | 4.41 | 20 | -42 | -10 |
| Decrease | PRC | L | 84 | 5.00 | -28 | -10 | -22 |
|  | PRC | R | 114 | 4.79 | 24 | -16 | -28 |
| 1:2:5 |  |  |  |  |  |  |  |
| Increase | Posterior HC/PHC | R | 12 | 5.12 | 18 | -42 | -4 |
| Decrease | PRC | L | 35 | 3.93 | -28 | -10 | -22 |
|  | PRC | R | 17 | 3.58 | 24 | -16 | -30 |
|  | PRC | R | 11 | 3.16 | 36 | -18 | -18 |
| 1:4:5 |  |  |  |  |  |  |  |
| Increase | - |  |  |  |  |  |  |
| Decrease | PRC | L | 102 | 5.37 | -28 | -6 | -22 |
|  | PRC | R | 143 | 4.98 | 24 | -16 | -24 |
|  | Anterior HC/PRC | R | 20 | 4.16 | 26 | 8 | -34 |

**Note.** H = hemisphere; *k* denotes cluster size; *x*, *y*, and *z* denote peak MNI coordinates; *t* is *t*-statistic. HC = hippocampus, PRC = perirhinal cortex, and PHC = parahippocampal cortex.

Critically, hippocampal sensitivity to increasing associative demands was observed when proximal pairs were placed closer to single items (1:2:5), and to a lesser extent, in the linear-spacing model (1:3:5). When proximal pairs were placed closer to distal pairs (1:4:5), no increases in hippocampal activity were observed. However, activity decreases in PRC were observed across all three models, with the strongest effect emerging under the association-biased model (1:4:5). Together, these findings indicate that neural sensitivity to associative demands is best captured when proximal pairs are treated as more similar to single item conditions, and that models weighting proximal pairs closer to distal conditions fail to account for hippocampal engagement.

#### 3.2.2 Between-condition contrasts

Supporting results from the parametric analyses, direct contrasts between encoding conditions revealed a clear dissociation along the anterior-posterior axis within the MTL. Overall, single item recollection was observed in more anterior regions compared to associative recollection which was localized more posteriorly. Specifically, for recollected hits, the single > distal contrast showed greater activity in left anterior hippocampus and right PRC. In contrast, the distal > single comparison revealed greater activation in bilateral posterior hippocampus. Contrasts involving the proximal condition further supported the idea that proximal pairs lie in an intermediate position along the item-to-association continuum, with no significant clusters found in either the proximal > single or proximal > distal contrasts. However, in line with the foregoing parametric analyses, the single > proximal yielded greater activation in right PHC, left anterior hippocampus/PRC, and left entorhinal cortex (ERC), while distal > proximal revealed robust activation in right posterior hippocampus.

#### 3.2.3 Exploratory brain-behavior correlation

To examine whether individual differences in MTL activity were associated with the behavioral memory advantage found for proximal relative to distal pairs, we conducted an exploratory brain-behavior regression analysis. Because the proximal > distal contrast directly indexed the behavioral effect of interest (better memory for proximal than distal pairs), mean contrast estimates from the anatomical hippocampus, PHC, and PRC ROIs were extracted for each participant^1^. Separate linear regression models were then fit for each ROI, with individual contrast estimates predicting proximal recollected d’, while controlling for distal recollected d’ to control for overall associative memory performance. Results indicated that PRC activity was significantly associated with proximal recollected d’ above and beyond general associative memory ability (β = 0.22, CI [0.04, 0.39], *t*(33) = 2.49, *p* = .018, adjusted *R²* = .19), suggesting that PRC engagement during successful proximal encoding contributes uniquely to the associative memory advantage for proximal pairs. No significant relationships were observed between proximal recollected d’ and HC or PHC mean contrast estimates (all *p*s > .39).

**Table 2.** Differences in univariate BOLD activity between conditions for recollected hits within the MTL.

| Contrasts | Regions | H | k | t | MNI Coordinates |  |  |
| --- | --- | --- | --- | --- | --- | --- | --- |
|  |  |  |  |  | x | y | z |
| Single > Distal | Anterior HC | L | 42 | 4.23 | -28 | -12 | -18 |
|  | Posterior PRC | R | 72 | 4.19 | 24 | -16 | -24 |
| Distal > Single | Posterior HC | R | 14 | 4.53 | 20 | -42 | -10 |
|  | Posterior HC | L | 9 | 4.27 | -24 | -42 | -10 |
| Single > Proximal | PHC | R | 59 | 4.45 | 30 | -6 | -24 |
|  | Anterior HC/PRC | L | 36 | 3.79 | -28 | -12 | -22 |
|  | ERC | L | 12 | 3.60 | -36 | 2 | -40 |
|  | PHC | R | 24 | 3.27 | 38 | 0 | -36 |
| Proximal > Single | - |  |  |  |  |  |  |
| Proximal > Distal | - |  |  |  |  |  |  |
| Distal > Proximal | Posterior HC | R | 20 | 7.49 | 18 | -42 | -6 |

**Note.** H = hemisphere; k denotes cluster size; x, y, and z denote peak MNI coordinates; t is t-statistic. HC = hippocampus, PRC = perirhinal cortex, and PHC = parahippocampal cortex.

**Figure 4.**
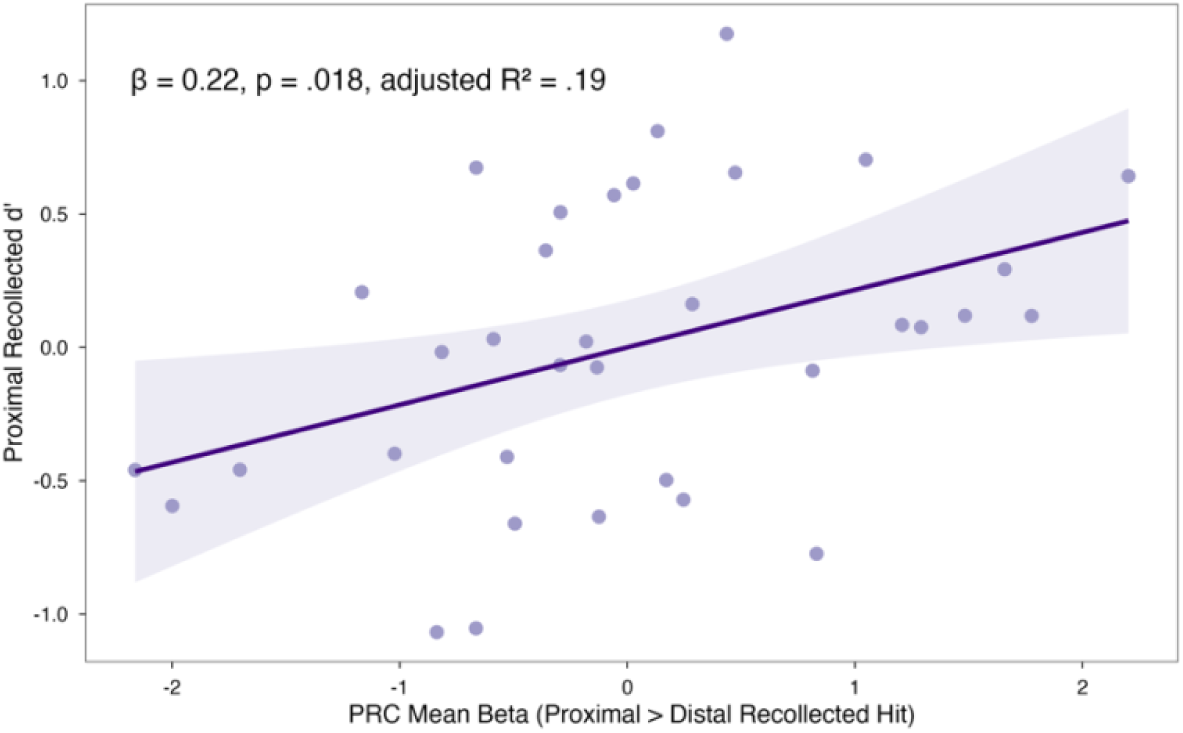
Partial regression plot showing the association between perirhinal cortex (PRC) mean beta estimate for the proximal > distal recollected hit contrast and proximal recollected d’, after controlling for distal recollected d’. Both axes represent residuals after regressing out distal recollected d’.

#### 3.2.4 MVPA classifiers

To assess neural discriminability between encoding conditions within MTL regions, we conducted ROI-based multivoxel pattern classification analyses for each pairwise comparison (single vs. distal, single vs. proximal, and proximal vs. distal). Classification accuracy was tested against chance performance (50%) using one-sample t-tests, with Benjamini-Hochberg correction across ROIs (hippocampus, PRC, and PHC).

Across all pairwise classification analyses, only the PHC consistently showed above-chance classification accuracy (see Figure 5). Specifically, PHC classification accuracy exceeded chance for single vs. distal (*M* = 0.58, *t*(34) = 4.40, *p* < .001), for single vs. proximal (*M* = 0.56, *t*(34) = 3.60, *p* = .002), and proximal vs. distal (*M* = 0.56, *t*(34) = 3.49, *p* = .002). The only other comparison showing above-chance classification accuracy was the hippocampus for single vs. distal (*M* = 0.53, *t*(34) = 2.00, *p* = .031).

**Figure 5.**
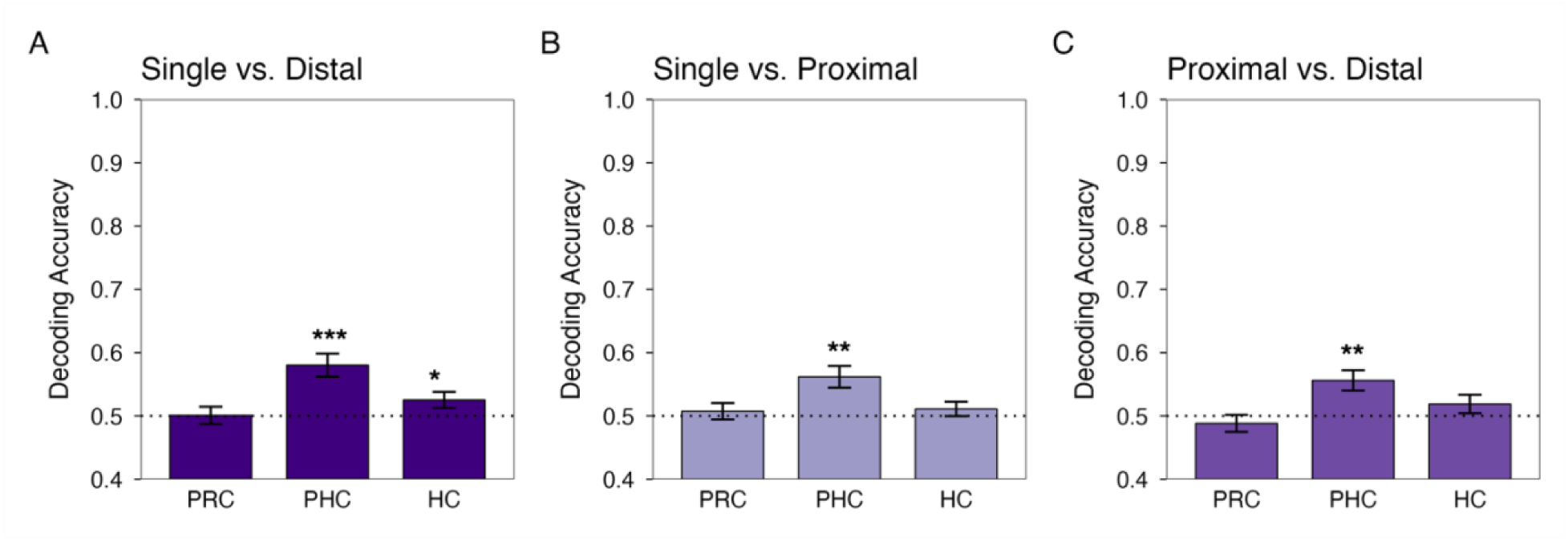
MTL subregion decoding accuracy for (A) single vs. distal, (B) single vs. proximal, (C) proximal vs. distal. Asterisks indicate significant pairwise comparisons (\**p* < .05, \*\**p* < .01, \*\*\**p* < .001). PRC = perirhinal cortex, and PHC = parahippocampal cortex, HC = hippocampus.

## 4 Discussion

The present study investigated whether spatial proximity supports associative memory in older adults and if so, how this benefit is reflected in MTL encoding activity. This study was motivated by previous work in the unitization field (Bridger et al., 2017; Carpenter & Dennis, 2023; Dennis et al., 2025; Huffer et al., 2022), and carried out by examining behavioral and neural responses to proximal (i.e., spatially integrated) arrangements of unrelated objects, in comparison to both distally arranged pairs of objects and to single objects. Behaviorally, older adults showed better memory for proximal compared to distal object pairs, though memory for proximal pairs remained below that of single items. Neurally, MTL activity varied as a function of associative demand, with increasing associative demand (distal > proximal > single) associated with increased activity in posterior hippocampus, whereas decreasing associative demand (distal < proximal < single) was associated with greater PRC activity. This pattern was more pronounced when proximal pairs were modeled as more closely resembling single items rather than distal pairs. Complementary to our parametric investigation of proximal pairs, between-condition contrasts revealed an anterior-posterior dissociation within the MTL, with single item recollection associated with robust anterior MTL activity including PRC and anterior hippocampus, and distal pair recollection associated with posterior hippocampus activity. Notably, the regions from the single > proximal contrast were largely the same as those from the single > distal contrast, and the regions from the distal > proximal contrast were largely the same as those from the distal > single contrast. Proximal pairs, however, did not show greater activity than either single items or distal pairs in any MTL region. This pattern suggests that proximal pairs shared neural properties with both single items and distal pairs, occupying an intermediate position with respect to MTL engagement. Moreover, the PHC reliably discriminated between encoding conditions across all binary comparisons. Exploratory brain-behavior correlation analyses further revealed that greater PRC engagement during proximal encoding uniquely predicted proximal memory above and beyond general associative memory ability. Together, these findings suggest that proximity-based unitization benefits associative memory in older adults by potentially shifting associative memory demands within the MTL, and that the structure of the aging MTL is preserved in a manner that situates proximal pairs at an intermediate position along the item-to-association continuum.

The behavioral memory advantage for proximal relative to distal image pairs is broadly consistent with prior unitization research demonstrating that perceptually integrating or grouping stimuli can facilitate associative memory performance (Bridger et al., 2017; Dennis et al., 2025; Huffer et al., 2022; Memel & Ryan, 2017). This benefit is particularly meaningful in the context of aging, as older adults are disproportionately impaired in associative memory (Naveh-Benjamin et al., 2004; Naveh-Benjamin & Mayr, 2018), an impairment attributed in part to reduced efficiency of hippocampal binding processes (for a review, see Giovanello & Dew, 2015).The finding that spatial proximity improved associative memory in older adults suggests that perceptual grouping can partially offset this age-related binding deficit, consistent with prior work showing that unitization benefits associative memory in older adults across various stimuli and contexts (Bastin et al., 2013; Huffer et al., 2022; Zheng et al., 2015, 2016). Notably, this benefit emerged without explicit unitization instructions, which is meaningful given that older adults are less likely than younger adults to spontaneously engage effective encoding strategies (Naveh-Benjamin et al., 2007), suggesting that perceptual grouping may support associative encoding in aging without requiring deliberate strategic effort.

Critically, like previously observed in a younger adult sample (Dennis et al., 2025), memory performance followed a graded pattern, with performance highest for single items, followed by proximal pairs, and lowest for distal pairs. Thus, although spatial proximity supported associative memory relative to distal pairs, proximal pairs continued to show poorer memory than single items. Prior studies demonstrating a memory benefit for unitized relative to non-unitized associations have often concluded that unitization effectively eliminates the associative memory deficit (Bridger et al., 2017; Huffer et al., 2022). However, these studies did not include a single item condition, and therefore could not directly evaluate whether unitized associations truly reached the level of item-based memory. The inclusion of a single item condition in the present study allowed a more direct test of this claim. The finding that memory performance for proximal pairs fell below single items suggests that while spatial proximity meaningfully reduced the associative memory burden, it did not eliminate the gap between associative and item-based memory, indicating that the associative deficit was attenuated rather than eliminated. This pattern of behavioral results is consistent with a continuum account of unitization, in which unitized associations occupy an intermediate position between item and associative memory rather than achieving full item-like status (Parks & Yonelinas, 2015).

Importantly, we were also able to examine whether proximal pairs’ intermediate representational position was closer to that of the single item or distal-associative ends of the continuum. To do so, we employed a parametric modulation approach that tested three related yet competing coding schemes: a linear-spacing model (1:3:5), a single-biased model (1:2:5), and an association-biased model (1:4:5), where 1 and 5 represented single and distal-associative conditions respectively, and the proximal weighting varied across models. This approach was motivated by the fact that the representational position of proximal pairs along the item-to-association continuum is not known a priori, and that different theoretical assumptions about the nature of proximal encoding yield different neural predictions. By testing all three models, we were able to ask not only whether MTL activity scaled with associative demand, but under what assumptions about proximal encoding this scaling emerged.

Within the BIC framework, single item memory is posited to rely more heavily on PRC-mediated processing, while associative memory places greater demands on hippocampal binding (Diana et al., 2007; Ranganath, 2010). Consistent with this framework, PRC activity increased as associative demand decreased across all three models, indicating that item-based PRC engagement was sensitive to the position of proximal pairs along the item-to-association continuum. Hippocampal activity, in contrast, increased with associative demand under the linear-spacing (1:3:5) and single biased (1:2:5) models, but disappeared entirely under the association-biased model (1:4:5), which positioned proximal pairs as neurally similar to distal pairs. When the model assumed proximal pairs were distal-like, it failed to capture hippocampal engagement, suggesting this coding scheme did not adequately track associative demand.

The observation that MTL activity scaled in a demand-dependent manner, with hippocampal engagement increasing as relational binding demands increased and PRC engagement increasing as item-based processing demands increased, indicates that the functional dissociation of item and associative processing described by the BIC model was preserved in our older adult sample (Diana et al., 2007; Ranganath, 2010). This is broadly consistent with prior work showing comparable MTL activity in younger and older adults during associative memory encoding (Addis et al., 2014; Memel & Ryan, 2017; Seo & Dennis, 2026). At the same time, the asymmetry in our parametric results raises additional considerations in the context of aging. Whereas PRC scaling was robust across all three models, hippocampal scaling was relatively modest and emerged under models that positioned proximal pairs closer to single items (i.e., linear-spacing and item-based). One possibility is that this asymmetry reflects the representational position of proximal pairs, such that hippocampal engagement for proximal encoding more closely resembled that of single items than distal pairs. An alternative possibility is that it reflects aging-related reductions in hippocampal recruitment during associative encoding. Older adults sometimes show reduced hippocampal activation during associative encoding relative to younger adults (Daselaar et al., 2003; Dennis et al., 2007, 2008; Mitchell et al., 2000), which may make hippocampal scaling more difficult to detect and more sensitive to how intermediate conditions are weighted. Under this account, the pattern of hippocampal scaling may reflect both a genuine reduction in binding demands for proximal pairs and a generally attenuated hippocampal signal in aging. Notably, the robustness of the PRC effect across all three models indicates that reduced overall signal cannot fully account for the observed pattern, supporting the interpretation that the parametric results reflect meaningful differences in how proximal pairs were represented rather than a global reduction in MTL sensitivity.

The dissociation along the anterior posterior axis of the MTL was complemented by the direct between-condition contrasts. Single item recollection was associated with greater anterior MTL activity relative to both associative conditions, including the left anterior hippocampus and right PRC, consistent with the BIC model’s account of anterior MTL supporting item-level representations (Diana et al., 2007; Ranganath, 2010). In contrast, distal pair recollection was associated with greater bilateral posterior hippocampal activity relative to both single items and proximal pairs, consistent with the relational binding demands of encoding spatially separated object pairs (Kumaran & Maguire, 2005; Staresina & Davachi, 2008, 2010). Mirroring the parametric results, these findings reveal an anterior-to-posterior shift in MTL activity from item-based to associative encoding, with anterior regions preferentially supporting item-level representations and posterior hippocampus supporting associative binding. Importantly, like the parametric findings, this pattern of results in older adults supports that the functional distinction of item and associative processing within the MTL was maintained in our sample.

Contrasts involving proximal pairs further clarified this organization. The single > proximal contrast revealed greater activity in left anterior hippocampus/PRC, a pattern that overlapped substantially with the single > distal contrast, which similarly revealed greater activity in left anterior hippocampus and right PRC. This overlap indicates that anterior hippocampus and PRC recruitment was largely unique to the single item condition relative to both associative conditions, supporting the interpretation that anterior MTL engagement was preferentially driven by item-based encoding processes. Thus, while proximal pairs showed a memory advantage over distal pairs, their encoding was not fully item-like with respect to PRC recruitment, as single items uniquely engaged anterior MTL beyond what was observed for proximal pairs. This pattern also indicates that proximal encoding continued to involve associative processing that was not captured by the item-based PRC activity observed for single items. Accordingly, this suggests that the PRC in older adults continues to support item-level processing by and large, rather than being recruited to support integrated associative representations, indicating that the item-specific processing of the PRC is maintained in aging.

With respect to associative processing, the distal > proximal contrast revealed greater right posterior hippocampal activity, and this pattern overlapped substantially with distal > single, which similarly showed greater bilateral posterior hippocampal activity. This indicates that posterior hippocampal recruitment was driven specifically by the distal condition relative to both single items and proximal pairs, suggesting that the relational binding load was greatest for spatially separated pairs. Correspondingly, proximal pairs did not require the degree of posterior hippocampal engagement observed for distal pairs, consistent with the interpretation that spatial proximity reduced, though did not eliminate, associative binding demands during encoding (Carpenter & Dennis, 2023; Dennis et al., 2024; Parks & Yonelinas, 2015).

Notably, unlike single items and distal pairs, each of which exhibited unique MTL activity relative to the other conditions, proximal pairs did not show greater MTL recruitment than either single items or distal pairs in any MTL subregion. This absence of unique proximal activity converges with the parametric modulation results in placing proximal pairs within the item-to-association continuum, sharing neural properties with both conditions without uniquely exceeding either. This suggests that spatial proximity did not recruit an additional or distinct MTL mechanism, but rather modulated the degree to which existing item- and association-based MTL processes were engaged. In aging, proximal pairs appeared to recruit the same anterior and posterior MTL systems as single items and distal pairs, but to an intermediate extent, consistent with the view that proximity supports associative memory by attenuating relational binding demands rather than by transforming associative information into a single item-like representation (Dennis et al., 2024; Parks & Yonelinas, 2015).

Importantly, our findings indicate that representational specificity within the MTL is retained in older adults. The anterior-posterior dissociation between item and associative encoding was robust, with single items and distal pairs each recruiting distinct MTL subregions. Classification analyses supported this specificity, demonstrating that hippocampal patterns reliably discriminated item from associative encoding, and that PHC patterns reliably discriminated all three conditions. Together, these findings indicate that proximal encoding engaged distinct item- and association-based processes. This representation aligns with prior findings in younger adults using this same paradigm, in which proximal pairs likewise occupied an intermediate representational position within the MTL, with proximal patterns more closely resembling those of distal pairs than single items (Dennis et al., 2025). The finding that proximal pairs do not achieve fully item-like status across age groups suggests that this intermediate position is a property of proximity-based unitization itself, rather than a consequence of aging.

Following up on univariate differences across MTL subregions, we investigated whether unitization through proximity resulted in differential neural patterns across encoding conditions. MVPA classification results revealed that the hippocampus showed above-chance classification for single compared to distal pairs, but not for comparisons involving proximal pairs. This is notable given the hippocampus’s established role in associative binding (Kumaran & Maguire, 2005; Olsen et al., 2012; Ranganath, 2010), a process that is disproportionately vulnerable in aging and thought to underlie age-related deficits in associative memory (for a review, see Giovanello & Dew, 2015). Despite well-documented age-related declines in associative memory performance, the current results show that the hippocampus continues to represent item and associative encoding in distinguishable ways. The results indicate that the hippocampal item-associative distinction is maintained in aging at the level of neural representation. However, at the same time, hippocampal patterns could not reliably differentiate proximal pairs from either single items or distal pairs. Like the foregoing univariate results, the multivariate results suggest that proximal encoding did not generate a hippocampal representation clearly distinguishable from either endpoint, placing proximal encoding in between item and associative memory processing (Carpenter & Dennis, 2023; Dennis et al., 2024; Parks & Yonelinas, 2015). Notably, a similar pattern was observed in younger adults, where hippocampal pattern similarity did not significantly differentiate proximal representations at encoding (Dennis et al., 2025), suggesting that the ambiguity of proximal representations within the hippocampus reflects their intermediate position along the item-to-association continuum rather than a feature of aging.

PHC was the only MTL region to show above-chance classification across all three pairwise comparisons. These results indicate that each encoding condition was represented in a systematically different pattern within PHC. This finding is consistent with PHC’s proposed role in processing contextual, spatial, and configurational properties of associative representations (Aminoff et al., 2007; Aminoff et al., 2013; Diana et al., 2007; Li et al., 2016; Staresina et al., 2011). For example, Aminoff et al. (2013) proposed that the PHC processes contextual associations that define meaningful environments and relationships among objects, and that PHC activity reflects sensitivity to relations among items and their configurational structure. In the current paradigm, the three encoding conditions differed systematically in their spatial arrangement and perceptual organization: single items involved no spatial pairing, proximal pairs involved spatially integrated objects, and distal pairs involved spatially separated objects. The PHC classifications likely reflected sensitivity to these configurational differences (Aminoff et al., 2013; Mullally & Maguire, 2011). The preservation of condition-specific PHC representations in our sample suggests that older adults remained sensitive to the spatial and configural properties that distinguish proximal from distal encoding, consistent with prior work suggesting intact context processing in older adults (Burke et al., 2018; Memel & Ryan, 2017; Robin & Moscovitch, 2017).

In contrast, the PRC did not show above-chance classification for any pairwise comparison. This may indicate that PRC represents items in a similar manner irrespective of the number of items in a display. This group-level null result stands in contrast to our exploratory brain-behavior analyses. Specifically, older adults who showed greater PRC activity for proximal relative to distal recollected pairs also demonstrated better proximal memory performance, even after controlling for overall associative memory ability. Together, these findings suggest that while the PRC may not encode conditions differently at the group level, the degree to which individual older adults recruited PRC during proximal encoding predicted their ability to form more integrated representations of proximal pairs. This interpretation is consistent with prior work implicating PRC in the encoding of integrated or holistically processed stimulus pairs (Ford et al., 2010; Haskins et al., 2008; Memel & Ryan, 2017), and with Parks and Yonelinas (2015), who proposed that greater PRC engagement reflects stronger associative integration. Although exploratory, this result raises the possibility that some older adults engage PRC-supported integrative processing more effectively during proximal encoding, contributing to individual variability in the proximal memory benefit. Such individual differences may relate to variability in PRC structure (Delhaye et al., 2019) as well as broader cognitive resource factors known to moderate associative memory benefits in aging, including working memory capacity and processing speed (Bouazzaoui et al., 2010; Murray & Kensinger, 2013; Naveh-Benjamin et al., 2007). Future work should examine whether such factors predict the neural mechanisms underlying proximity-based associative encoding (Becker et al., 2015; Carr et al., 2017; Zheng et al., 2017).

The foregoing results should be considered in the context of the following study limitations. First, the proximal condition manipulated both spatial proximity and logical orientation, such that proximal pairs were spatially integrated and configured in a meaningful arrangement. Importantly, the degree of configural meaning likely varied across pairs. For example, an otter and cleaver configured with the cleaver’s handle touching the otter’s paw may form a more coherent unit compared to a door lock resting atop a bathtub. That is, even though both pairs went through the same norming process to have the most logical proximal configuration between their constituent objects, variance in the action affordances of those objects (e.g., an otter paw’s capacity to grasp, a cleaver’s capacity to be wielded) may have contributed to variance in configural meaning between proximal pairs. Because spatial proximity and configural meaning were not varied independently, their relative contributions to the observed behavioral and neural effects cannot be isolated, and future work systematically dissociating these dimensions would help clarify which aspects of proximity support associative memory in aging. Second, the brain-behavior analyses reported here were exploratory and based on a modest sample; these findings should be interpreted with caution and replicated in larger samples.

## 5 Conclusion

In sum, the present findings suggest that proximity-based unitization benefits associative memory in older adults by shifting how associative information is processed within the MTL. Proximal pairs occupied an intermediate position within the anterior-posterior MTL encoding gradient, sharing anterior MTL properties with single items and posterior hippocampal properties with distal pairs, without requiring unique MTL recruitment beyond either condition. This pattern, consistent across parametric modulation, between-condition contrast, and multivariate analyses, indicates that spatial proximity modulated the degree to which existing item- and association-based MTL processes were engaged, rather than producing a complete shift toward item-level processing (Dennis et al., 2025; Parks and Yonelinas, 2015).

Critically, these findings speak to the flexibility of MTL function in aging. Encoding activity in older adults aligned with the functional organization described by the BIC model (Diana et al., 2007; Ranganath, 2010). Critically, however, this organization was not fixed. A simple manipulation of object placement at encoding was sufficient to shift MTL processing in older adults, such that proximal pairs engaged item- and association-based MTL systems to an intermediate degree. The fact that this shift occurred in the absence of any explicit instruction to unitize indicates that the aging MTL retains the capacity to modulate its processing in response to the configural structure of incoming information (Dennis et al., 2019; Memel & Ryan, 2017).

Exploratory brain-behavior analyses further revealed that individual differences in PRC engagement were associated with proximal memory performance above and beyond general associative memory ability. As such, results suggest that variability in integrative encoding may be an important determinant of who benefits most from proximity in aging. Together, these findings highlight the potential of incidental perceptual grouping to support associative memory in aging, and underscore the value of directly comparing unitized associations against both item and associative conditions to characterize the nature of integrative encoding within the aging MTL.

## 6 CRediT author statement

**Min Sung Seo:** Conceptualization, Methodology, Software, Validation, Formal Analysis, Data Curation, Writing – Original Draft, Writing – Review & Editing, Visualization, Project administration; **Alexa Becker:** Investigation, Data Curation, Writing – Review & Editing; **Lin-Han Hannah Huang:** Conceptualization, Writing – Original Draft, Writing – Review & Editing, Visualization; **Amy A. Overman:** Writing – Review & Editing, Funding acquisition; **Nancy A. Dennis:** Conceptualization, Methodology, Resources, Writing – Review & Editing, Supervision, Project administration, Funding acquisition

## 8 Acknowledgements

The authors acknowledge use of the Social, Life, and Engineering Sciences Imaging Center (SLEIC), Pennsylvania State University, University Park, PA (RRID:SCR_014922). We acknowledge the use of the Human MRI Facility for MRI data collection, experimental consultation, and technical support.

## Footnotes

1 Although the proximal > distal contrast did not yield suprathreshold clusters at the group-level, group-level null effects do not preclude meaningful between-subject variability in neural responses

